# High virulence does not necessarily impede viral adaptation to a new host: A case study using a plant RNA virus

**DOI:** 10.1101/060137

**Authors:** Anouk Willemsen, Mark P. Zwart, Santiago F. Elena

**Affiliations:** Instituto de BiologÍa Molecular y Celular de Plantas (IBMCP), Consejo Superior de Investigaciones CientÍficas-Universidad Politècnica de Valencia, Campus UPV CPI 8E, Ingeniero Fausto Elio s/n, 46022 València, Spain.; Institute of Theoretical Physics, University of Cologne, Zülpicher Straße 77, 50937 Cologne, Germany.; The Santa Fe Institute, 1399 Hyde Park Road, Santa Fe, NM 87501, USA.

**Keywords:** Adaptation, Experimental evolution, Host-pathogen interactions, Virulence, Virus evolution, Genome architecture evolution

## Abstract

**Background:** When between-host selection pressures predominate, theory suggests that high virulence could hinder between-host transmission of microparasites, and that virulence therefore will evolve to lower levels that optimize between-host transmission. Highly virulent microparasites could also curtail host development, thereby limiting both the host resources available to them and their own within-host effective population size. High virulence might therefore curtail the mutation supply rate and increase the strength with which genetic drift acts on microparasite populations, thereby limiting the potential to adapt to the host and ultimately perhaps the ability to evolve lower virulence. As a first exploration of this hypothesis, we evolved *Tobacco etch virus* carrying an eGFP fluorescent marker in two semi-permissive host species, *Nicotiana benthamiana* and *Datura stramonium*, for which it has a large difference in virulence. We compared the results to those previously obtained in the typical host, *Nicotiana tabacum*, where we have shown that carriage of *eGFP* has a high fitness cost and its loss serves as a real-time indicator of adaptation.

**Results:** After over half a year of evolution, we sequenced the genomes of the evolved lineages and measured their fitness. During the evolution experiment, marker loss leading to viable virus variants was only observed in one lineage of the host for which the virus has low virulence, *D. stramonium*. This result was consistent with the observation that there was a fitness cost of *eGFP* in this host, while surprisingly no fitness cost was observed in the host for which the virus has high virulence, *N. benthamiana*. Furthermore, in both hosts we observed few lineages with increases in viral fitness, and host-specific convergent evolution at the genomic level was only found in *N. benthamiana*.

**Conclusions:** The results of this study do not lend support to the hypothesis that high virulence impedes microparasites’ evolution. Rather, they exemplify that jumps between host species can be game changers for evolutionary dynamics. When considering the evolution of genome architecture, host species jumps might play a very important role, by allowing evolutionary intermediates to be competitive.

## Background

From both applied and fundamental perspectives, virulence is a key phenotypic trait of microparasites. In medicine and agriculture, it is crucial to understand mechanistically how microparasites harm the host, in order to devise effective interventions. From a more fundamental perspective, evolutionary biologists have long been interested in understanding why many microparasites are highly virulent. It has been suggested that virulence reduces between-host transmission, and that selection would therefore act to maximize between-host transmission by reducing virulence [1, 2]. High virulence would signal maladaptation, for example following a host-species jump, and eventually be selected against. The ubiquity of microparasitic virulence and the fact that many apparently well-adapted microparasites have high virulence led to a more sophisticated framework: the hypothesis that there are tradeoffs between virulence and transmission [2–5]. This framework posits that high levels of replication could increase the probability of a microparasite being transferred to a new host, whilst also increasing the probability that the host would die quickly and the temporal window for transmission would be very brief. Under this more plausible framework, virulence evolves to the level that optimizes between-host transmission [4, 6, 7].

The tradeoff hypothesis forms the cornerstone for theoretical frameworks considering the evolution of virulence in many different pathosystems. Many important additions to the framework have been made, for example recognizing that within-host competition and opportunism can lead to increases in virulence [8–10]. Moreover, the importance of other factors at the between-host level have been given consideration, such as self-shading [11]. Self-shading occurs when the host population is structured, and a highly virulent microparasite kills all host organisms in a subpopulation before transmission to another subpopulation can be achieved. The effects of evolution on microparasitic virulence have therefore been given considerable attention, although the number of experimental studies that address this issue is still rather limited, especially for viruses [12].

The effects of evolution on microparasite virulence have been widely considered. However, virulence itself could also have profound effects on evolution, including its own evolutionary dynamics [13]. This reversed causality is already apparent from the tradeoff model, under which microparasites with suboptimal virulence will undergo reduced between-hosts transmission. All other things equal, if a smaller number of hosts are infected effective population size will be decreased, increasing the strength of genetic drift and decreasing the mutation supply rate. In addition, the evolution to optimum virulence may be slow as this optimum is not static and can shift towards lower virulence as the density of susceptible hosts decreases [14]. Moreover, a wide range of virulence can be associated with each step of evolution towards the optimum, where selection favors genotypes with higher fitness that may improve transmission but not necessarily improve virulence [13]. Besides these effects of virulence on evolution, it is conceivable that a similar within-host effect could also occur, when virulence curtails host development and thereby limits the host resources available to the microparasite. Virulence would then limit the microparasite effective population size within hosts, again reducing the mutation supply and thereby slowing the rate of adaptation. Interestingly, all of these mechanisms could limit the rate at which lower virulence evolves, meaning that high virulence might persist longer than suggested by the simple tradeoff model [13].

There are many reasons why high virulence in host-pathogen interactions could emerge, but the most likely avenue is probably a change of host species. For example, infection of Ebola virus in bats is asymptomatic, while in humans and other primates the death rate is high [15]. Changes in virulence have been explained by the host phylogeny, where similar levels of virulence are displayed by closely related hosts and host jumps across large genetic distances may result in high virulence [16]. However, if a microparasite is confronted with a new host environment in which its level of virulence is altered, how does virulence affect its ability to adapt to the new host?

Here we address this question using *Tobacco etch virus* (genus: *Potyvirus*, family: *Potyviridae*), a (+)ssRNA virus that infects a wide-range of host plants, and an experimental evolution approach. To consider the effect of virulence on virus adaptation, we looked for two natural host species in which (*i*) there was some evidence that TEV potential for adaptation would be roughly similar, and (*ii*) there was a large difference in virulence. The distribution of mutational fitness effects (DMFE) of TEV has been compared in eight host species, and this study concluded that there were strong virus genotype-by-host species interactions [17]. For many host species distantly related to the typical host of TEV, *Nicotiana tabacum*, the DMFE changed drastically; many mutations that were neutral or deleterious in *N. tabacum*, became beneficial. However, for two closely related host species, *Nicotiana benthamiana* and *Datura stramonium*, most mutations tested remained neutral or deleterious [17], implying that the fraction of beneficial mutations in both hosts is small. Moreover, virus accumulation after one week of infection is also similar for both hosts [18]. On the other hand, TEV infection of *N. benthamiana* will typically result in heavy stunting and the death of the plant within a matter of weeks, whereas TEV infection of *D. stramonium* is virtually asymptomatic. Whilst there are many similarities between TEV infection in these two hosts, one key difference is host-pathogen interactions and therewith levels of viral virulence brought about.

As a first exploration of the effects of virulence on microparasite evolution, we therefore decided to evolve TEV in *N. benthamiana* and *D. stramonium.* By serially passaging each independent lineage in a single plant, our study maximizes within-host selection. This setup allows us to exclusively focus on effects of within-host selection, although for our model system we expect to see large differences in the resulting population size and the scope of virus movement within the host. Moreover, to immediately gauge whether adaptive evolution might be occurring, we passaged a TEV variant expressing a marker protein (Fig. 1), the enhanced GFP (eGFP). Upon long-duration passages in *N. tabacum*, this exogenous sequence is quickly lost due to its strong fitness cost, and its loss is reliably indicated by a loss of eGFP fluorescence [19]. According to the above hypothesis that high virulence may impair the rate of microparasite evolution, we expect that adaptive evolution would occur more quickly in the host species for which TEV has lower virulence, *D. stramonium*, than in the host species for which it has high virulence, *N. benthamiana*. Hence, we expected that in *D. stramonium* (*i*) the eGFP marker would be lost more rapidly, (*ii*) there would be more sequence-level convergent evolution, and (*iii*) there would be larger increases in within-host competitive fitness. However, the results clashed with these simple hypotheses, exemplifying the extent to which a host species jump can be a game changer for RNA virus evolutionary dynamics.

**Figure 1.**
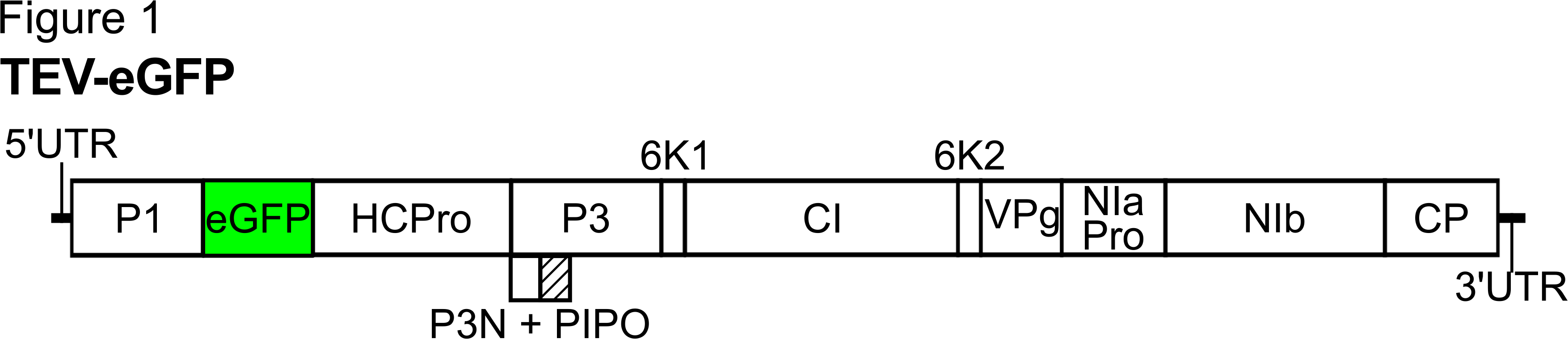
Schematic representation of TEV-eGFP. The *eGFP* gene is located between *P1* and *HC-Pro* cistrons. Proteolytic cleavage sites were provided at both ends of *eGFP.*

## Methods

### Virus stocks, plants and serial passages

The TEV genome used to generate TEV-eGFP virus, was originally isolated from *N. tabacum* plants [20]. To generate a virus stock of the ancestral TEV-eGFP, the pMTEV-eGFP plasmid [21] was linearized by digestion with *Bgl*II prior to *in vitro* RNA synthesis using the mMESSAGE mMACHINE^®^ SP6 Transciption Kit (Ambion), as described in [22]. The third true leaf of 4-week-old *N. tabacum* L var Xanthi *NN* plants was mechanically inoculated with 5 μg of transcribed RNA. All symptomatic tissue was collected 7 days post-inoculation (dpi). For the serial passage experiments, 500 μg homogenized stock tissue was ground into fine powder and diluted in 500 μl phosphate buffer (50 mM KH_2_PO_4_, pH 7.0, 3% polyethylene glycol 6000). From this mixture, 20 μl were then mechanically inoculated on the sixth true leaf of 4-week old *N. benthamiana* Domin plants and on the third true leaf of 4-week old *D. stramonium* L plants. Ten independent replicates were used for each host plant. Based on a previous study done in *N. tabacum* [19], passages of TEV-eGFP in *D. stramonium* were done every 9 weeks. In *N. benthamiana* the virus induces host mortality, and therefore the passages had to be restricted to 6 weeks for this host. At the end of the designated passage duration all leaves above the inoculated one were collected and stored at −80 °C. For subsequent passages the frozen tissue was homogenized and a sample was ground and resuspended with an equal amount of phosphate buffer [19]. Then, new plants were mechanically inoculated as described above. Three passages were performed for lineages evolved in *D. stramonium* and five passages for lineages evolving in *N. benthamiana*, making the number of generations of evolution similar in both hosts. All plants were kept in a biosafety level 2 greenhouse at 24° C with 16 h light:8 h dark photoperiod.

### Reverse transcription polymerase chain reaction (RT-PCR)

To determine whether deletions occurred at the *eGFP* locus, RNA was extracted from 100 mg homogenized infected tissue using the InviTrap Spin Plant RNA Mini Kit (Stratec Molecular). Reverse transcription (RT) was performed using M-MuLV reverse transcriptase (Thermo Scientific) and reverse primer 5’-CGCACTACATAGGAGAATTAG-3’ located in the 3’UTR of the TEV genome. PCR was then performed with Taq DNA polymerase (Roche) and primers flanking the *eGFP* gene: forward 5’-GCAATCAAGCATTCTACTTC-3’, and reverse 5’-CCTGATATGTTTCCTGATAAC-3’. PCR products were resolved by electrophoresis on 1% agarose gels.

### Virus accumulation and within-host competitive fitness assays

Prior to performing assays, the genome equivalents per 100 mg of tissue of the ancestral virus stocks and all evolved lineages were determined for subsequent fitness assays. The InviTrap Spin Plant RNA Mini Kit (Stratec Molecular) was used to isolate total RNA of 100 mg homogenized infected tissue. Real-time quantitative RT-PCR (RT-qPCR) was performed using the One Step SYBR PrimeScript RT-PCR Kit II (Takara), in accordance with manufacturer instructions, in a StepOnePlus Real-Time PCR System (Applied Biosystems). Specific primers for the *CP* gene were used: forward 5’-TTGGTCTTGATGGCAACGTG-3’ and reverse 5’-TGTGCCGTTCAGTGTCTTCCT-3’. The StepOne Software v.2.2.2 (Applied Biosystems) was used to analyze the data. The concentration of genome equivalents per 100 mg of tissue was then normalized to that of the sample with the lowest concentration, using phosphate buffer.

For the accumulation assays, 4-week-old *N. benthamiana* and *D. stramonium* plants were mechanically inoculated with 50 μl of the normalized dilutions of ground tissue. Inoculation of each viral lineage was done on the same host plant on which it had been evolved, plus TEV and the ancestral TEV-eGFP virus on each of the hosts, using three independent plant replicates per lineage. Leaf tissue was harvested 10 dpi. Total RNA was extracted from 100 mg of homogenized tissue. Virus accumulation was then determined by means of RT-qPCR for the *CP* gene of the ancestral and the evolved lineages. For each of the harvested plants, at least three technical replicates were used for RT-qPCR.

To measure within-host competitive fitness, we used TEV carrying a red fluorescent protein: TEV-mCherry as a common competitor. This virus has a similar insert size and within-host fitness compared with TEV-eGFP [19]. All ancestral and evolved viral lineages were again normalized to the sample with the lowest concentration, and 1:1 mixtures of viral genome equivalents were made with TEV-mCherry [21]. The mixture was mechanically inoculated on the same host plant on which it had been evolved, plus TEV and the ancestral TEV-eGFP virus on each of the hosts, using three independent plant replicates per viral lineage. The plant leaves were collected at 10 dpi, and stored at −80 °C. Total RNA was extracted from 100 mg homogenized tissue. RT-qPCR for the *CP* gene was used to determine total viral accumulation, and independent RT-qPCR reactions were also performed for the mCherry sequence using specific primers: forward 5’-CGGCGAGTTCATCTACAAGG-3’ and reverse 5’-TGGTCTTCTTCTGCATTACGG-3’. The ratio of the evolved and ancestral lineages to TEV-mCherry (*R*) is then *R* = (*n_CP_* − *n_mcherry_*)/*n_mcherry_*, where and are the RT-qPCR measured copy numbers of *n_CP_* and *n_mCherry_*, respectively. Then we can estimate the within-host competitive fitness as 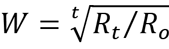 where *R_0_* is the ratio at the start of the experiment and *R_t_* the ratio after *t* days of competition [22]. The statistical analyses comparing the fitness between lineages were performed using R v.3.2.2 [23] and IBM SPSS Statistics version 23.

### Illumina sequencing, variants, and SNP calling

For Illumina next-generation sequencing (NGS) of the evolved and ancestral lineages, the viral genomes were amplified by RT-PCR using AccuScript Hi-Fi(Agilent Technologies) reverse transcriptase and Phusion DNA polymerase (Thermo Scientific), with six independent replicates that were pooled. Each virus was amplified using three primer sets, generating three amplicons of similar size (set 1: 5’-GCAATCAAGCATTCTACTTCTATTGCAGC-3’ and 5’-CCTGATATGTTTCCTGATAAC-3’; set 2: 5’-ACACGTACTGGCTGTCAGCG-3’ and 5’-CATCAATGTCAATGGTTACAC-3’; set 3: 5’-CCCGTGAAACTCAAGATAG-3’ and 5’-CGCACTACATAGGAGAATTAG-3’). Equimolar mixtures of the three PCR products were made. Sequencing was performed at GenoScreen (Lille, France: http://www.genoscreen.com). Illumina HiSeq2500 2×100bp paired-end libraries with dual-index adaptors were prepared along with an internal PhiX control. Libraries were prepared using the Nextera XT DNA Library Preparation Kit (Illumina Inc.). Sequencing quality control was performed by GenoScreen, based on PhiX error rate and Q30 values.

Read artifact filtering and quality trimming (3’ minimum Q28 and minimum read length of 50 bp) was done using FASTX-Toolkit v.0.0.14 [24]. De-replication of the reads and 5’ quality trimming requiring a minimum of Q28 was done using PRINSEQ-lite v.0.20.4 [25]. Reads containing undefined nucleotides (N) were discarded. Initially, the ancestral TEV-eGFP sequence was mapped using Bowtie v.2.2.6 [26] against the reference TEV-eGFP sequence (GenBank accession: KC918545). Error correction was done using Polisher v2.0.8 (available for academic use from the Joint Genome Institute) and a consensus sequences was defined for the ancestral TEV-eGFP lineage. Subsequently, the cleaned reads of the evolved sequences were mapped using Bowtie v.2.2.6 against the new defined consensus sequence. Single nucleotide mutations for each viral lineage were identified using SAMtools’ mpileup [27] and VarScan v.2.3.9 [28], where the maximum coverage was set to 40000 and mutations with a frequency < 1% were discarded. Note that the single nucleotide mutations detected here can be fixed (frequency > 50%) in the evolved lineages, as the detection was done over the ancestral population. Hence, it allows us to compare the mutations that arose by evolving TEV-eGFP in the different hosts.

## Results

### Experimental setup and fluorescent marker stability upon passaging of TEV-eGFP

TEV-eGFP was mechanically passaged in *N. benthamiana* and *D. stramonium*. In a previous study we noted that 9-week long passages led to rapid deletion of *eGFP* as well as rapid convergent evolution in *N. tabacum* [19]. Although 9-week passages could be performed in *D. stramonium*, for *N. benthamiana* this was not possible due to virus-induced host mortality. These plants died after 6 weeks of infection, and therefore we were forced to collect tissue at this time point. As *D. stramonium* grows to similar heights as *N. tabacum* when infected with TEV, and *N. benthamiana* does not grow much after infection, we chose to maximize infection duration to make the results comparable to those obtained in *N. tabacum* [19]. We performed three 9-week passages in *D. stramonium* and – to keep the total evolutionary time comparable – five 6-week passages in *N. benthamiana*. In *D. stramonium* all ten lineages initiated were completed, whereas in *N. benthamiana* only 6/10 lineages were completed. The remaining four *N. benthamiana* lineages failed to cause infection in subsequent rounds of passaging, and were therefore halted. Initial symptomatology of TEV-eGFP in *N. benthamiana* was very mild, while this symptomatology was more severe in the second and subsequent passages, possibly indicating adaptation of the virus to this alternative host. In *D. stramonium* the symptomatology was constant along the evolution experiment.

Based on previous results, we expected that the exogenous *eGFP* gene sequence would be rapidly purged [19, 29, 30], and as such would serve as a first indicator of the occurrence of TEV adaptation. However, the usefulness of fluorescence for determining the integrity of the *eGFP* marker was limited in both hosts, by (i) the high levels of autofluorescence in the highly symptomatic *N. benthamiana* leaves, and (ii) the patchy fluorescence in the *D. stramonium* tissue. Therefore, unlike for TEV-eGFP in *N. tabacum,* the fluorescent marker was of limited use here. Nevertheless, all *N. benthamiana* lineages appeared to have some fluorescence until the end of the evolution experiment, and we observed a loss of fluorescence in only 1/10 *D. stramonium* lineages in the third 9-week passage.

After each passage, RNA was extracted from the collected leaf tissue, and RT-PCR with primers flanking the *eGFP* insert was performed. This RT-PCR assay can therefore detect deletions in the *eGFP* gene, even when deletions extend well into the downstream HC-Pro cistron [19]. In general, the RT-PCR results confirmed the fluorescence microscopy results: A large deletion was detected only in the one *D. stramonium* lineage with a loss of fluorescence (Fig. 2A; 9-weeks passage 2 L8). This deletion variant went to a high frequency in the subsequent passage (Fig. 2A; 9-weeks passage 3 L8). For *N. benthamiana* lineages, we did detect a low-frequency deletion in the *eGFP* cistron in one lineage (Fig. 2B; 6-weeks passage 4 and 5 L4), but this deletion is so large that this variant is most likely no longer capable of autonomous replication. The deletion size is around 1500 nt, which means that after deleting the entire *eGFP,* around 800 nt are deleted from HC-Pro, which has a size of 1377 nt in total. This deletion extends well into the central region of HC-Pro, beyond the well-conserved FRNK box, which is essential for virus movement and RNA-silencing suppressor activity [31, 32]. We performed an extra round of passaging with all *N. benthamiana* lineages to check whether this variant would remain at a low frequency, and found exactly this result (Fig. 2B]; 6-week passage 6 L4). Furthermore, we detected a small deletion in one lineage (Fig. 2B; 6-week passage 5 and 6 L1) that was maintained at a low frequency in subsequent passages of the virus population.

**Figure 2.**
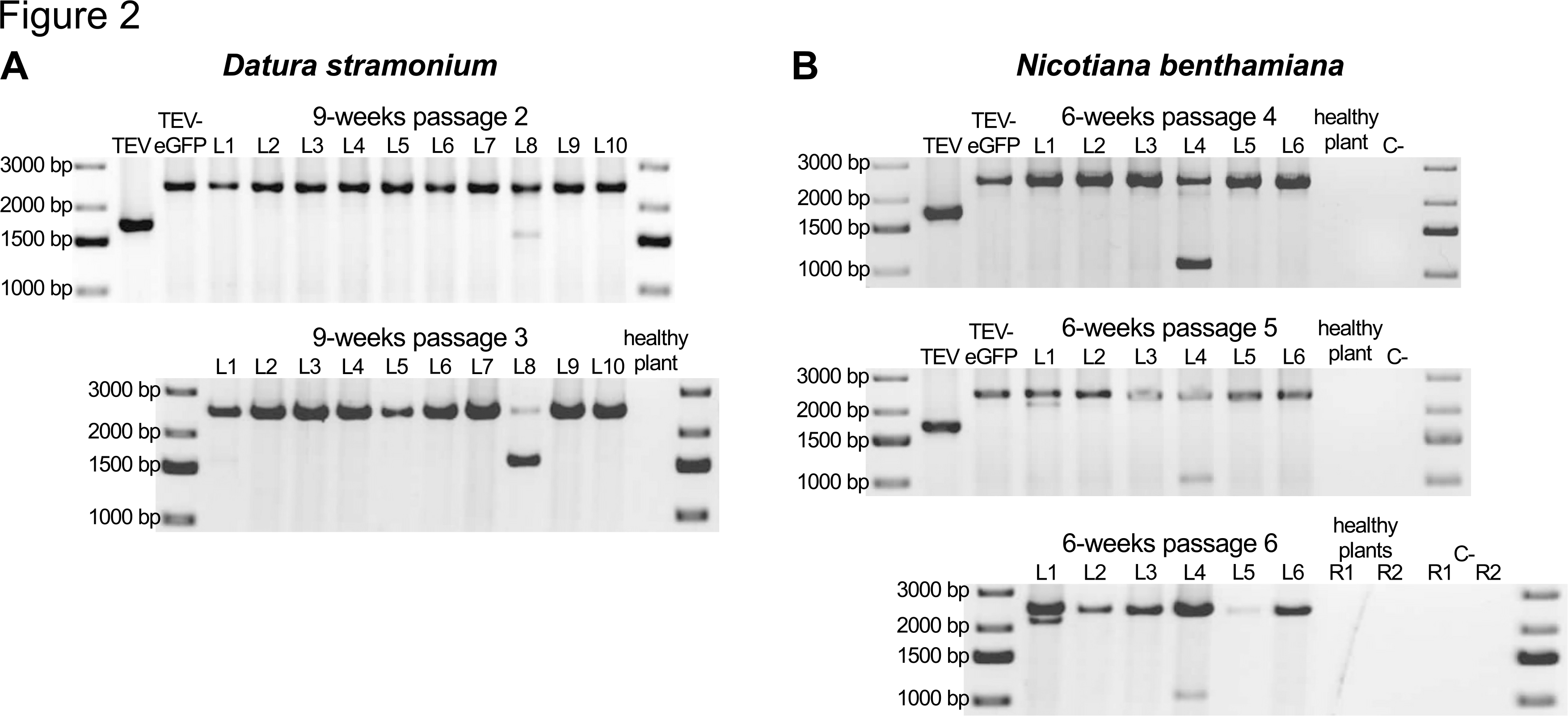
Deletion detection in the *eGFP* gene. Agarose gels with RT-PCR products of the region encompassing the *eGFP* gene. The TEV and TEV-eGFP are shown for comparative purposes. The negative controls are healthy plants and PCR controls (C-). (A) TEV-eGFP in *D. stramonium* has 10 independent lineages (L1-L10). A deletion encompassing the *eGFP* gene was detected in one lineage (L8) in the second 9-week passage. This deletion went to a high frequency in the subsequent passage. (B) TEV-eGFP in *N. benthamiana* has six independent lineages (L1-L6). A deletion bigger than the size of *eGFP* was detected in one lineage (L4) in the fourth 6-week passage. This deletion was not fixed in the two subsequent passages. A small deletion was detected in the fifth and sixth 6-week passage in L1.

### Whole-genome sequencing of the evolved lineages

All evolved and the ancestral TEV-eGFP lineages were fully sequenced by Illumina technology (SRA accession: SRP075180). The consensus sequence of the ancestral TEV-eGFP population was used as a reference for mapping the evolved lineages. The deletion observed by RT-PCR (Fig. 2A) in one of the *D. stramonium* lineages was confirmed by a low number of reads mapping inside the *eGFP* region (median coverage *eGFP:* 111.5), compared to a higher average coverage outside this region (median coverage *P1* gene: 19190, median overall genome coverage: 18460). The large deletion included the N-terminal region of *HC-Pro,* as observed for other deletions that occur after gene insertions before this gene [19, 34]. For all other lineages in *D. stramonium* and *N. benthamiana,* coverage over the genome was largely uniform and similar to the ancestral virus population, indicating that there were indeed no genomic deletions present at appreciable frequencies.

Single nucleotide mutations were detected from a frequency as low as 1%, comparing the evolved TEV-eGFP lineages in *N. benthamiana* and *D. stramonium* to the ancestral population (Fig. 3). This detection was also performed for evolved TEV-eGFP lineages in *N. tabacum,* that were sequenced in a previous study [19] (SRA accession: SRP075228). In the evolved *N. benthamiana* lineages 165 unique mutations were found, with a median of 34.5 (range: 27 - 47) mutations per lineage. In the evolved *D. stramonium* lineages 239 unique mutations were found, with a median of 31.5 (range: 16 - 35) mutations per lineage. In the evolved *N. tabacum* lineages, 183 unique mutations were found, with a median of 21.5 (range: 17 - 36) mutations per lineage. Note that the single nucleotide mutations detected here can be fixed (frequency > 50%) in the evolved lineages, as the detection was done over the ancestral population. Hence, it allows us to compare the mutations that arose by evolving TEV-eGFP in the different hosts.

**Figure 3.**
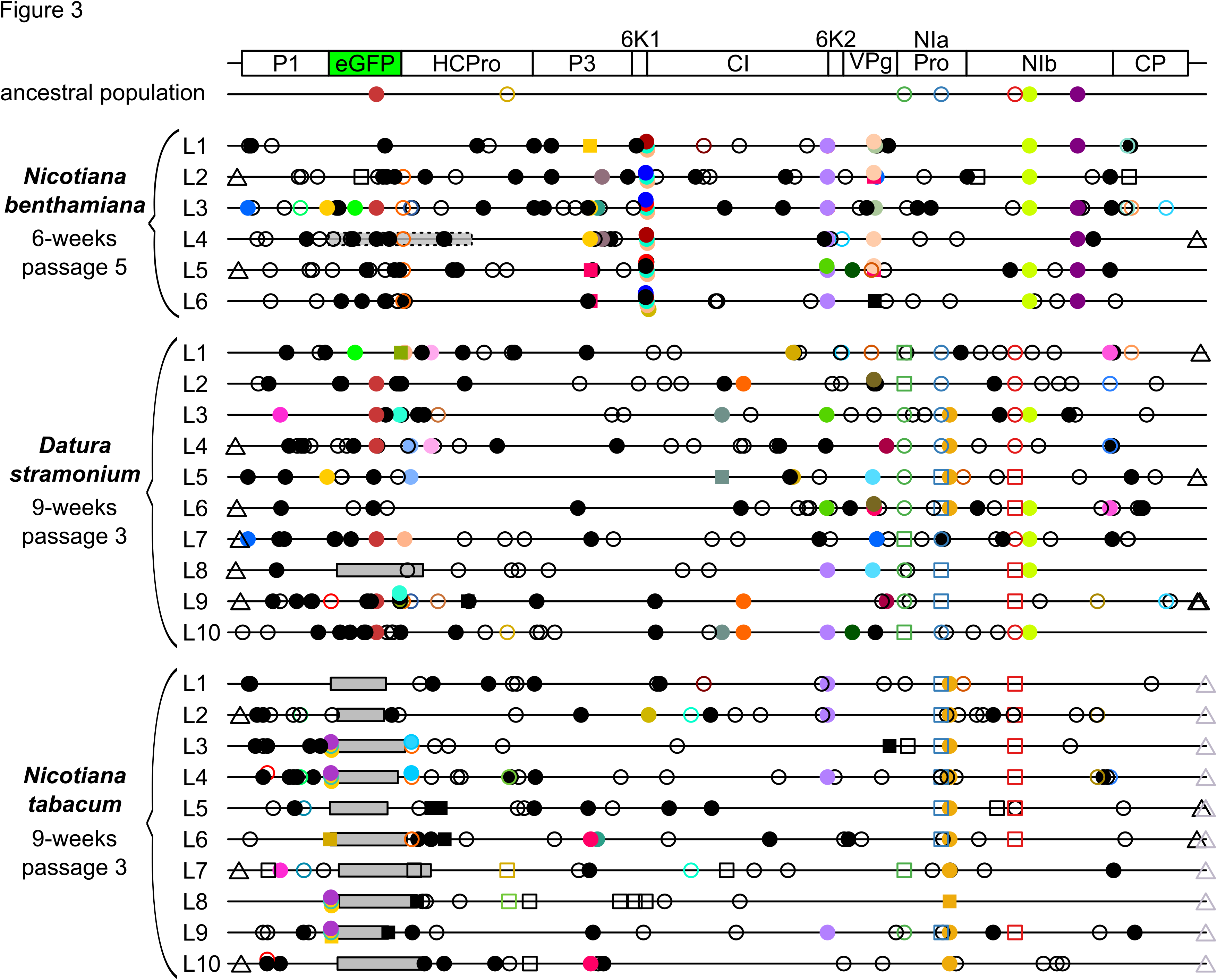
Genomes of the TEV-eGFP lineages evolved the three different hosts as compared to the ancestral lineage. Mutations were detected using NGS data of the evolved lineages (L1-L10), as compared to their ancestral population. The square symbols represent mutations that are fixed (> 50%) and the circle symbols represent mutations that are not fixed (< 50%). Filled symbols represent nonsynonymous substitutions and open symbols represent synonymous substitutions. The triangle symbols represent mutations that are present in either the 3‵UTR or 5‵UTR. Black substitutions occur only in one lineage, whereas color-coded substitutions are repeated in two or more lineages. Note that the mutations are present at different frequencies as reported by VarScan 2. Grey boxes with continuous black lines indicate genomic deletions in the majority variant of the virus population. The grey transparent box with dotted black lines in L4 of *N. benthamiana* indicates a genomic deletion in a minority variant. The latter box was drawn to indicate the size of the deletion, assuming that the deletion starts at the first position of *eGFP.* For more information on the frequency of the mutations please see Additional file 2: Tables S1-S3.

We detected only one mutation (U6286C; CI/Y2096H) that is shared between all three hosts. However, this mutation was present at a low frequency and not detected in all *D. stramonium* and *N. tabacum* lineages (Fig. 3 and Table 1). The *N. benthamiana* and *D. stramonium* lineages share more mutations (15) than either *N. benthamiana* or *D. stramonium* share with *N. tabacum* (4 and 9, respectively). However, most of these mutations are present in only a few lineages and at low frequency (Fig. 3 and Table 1).

**Table 1.**
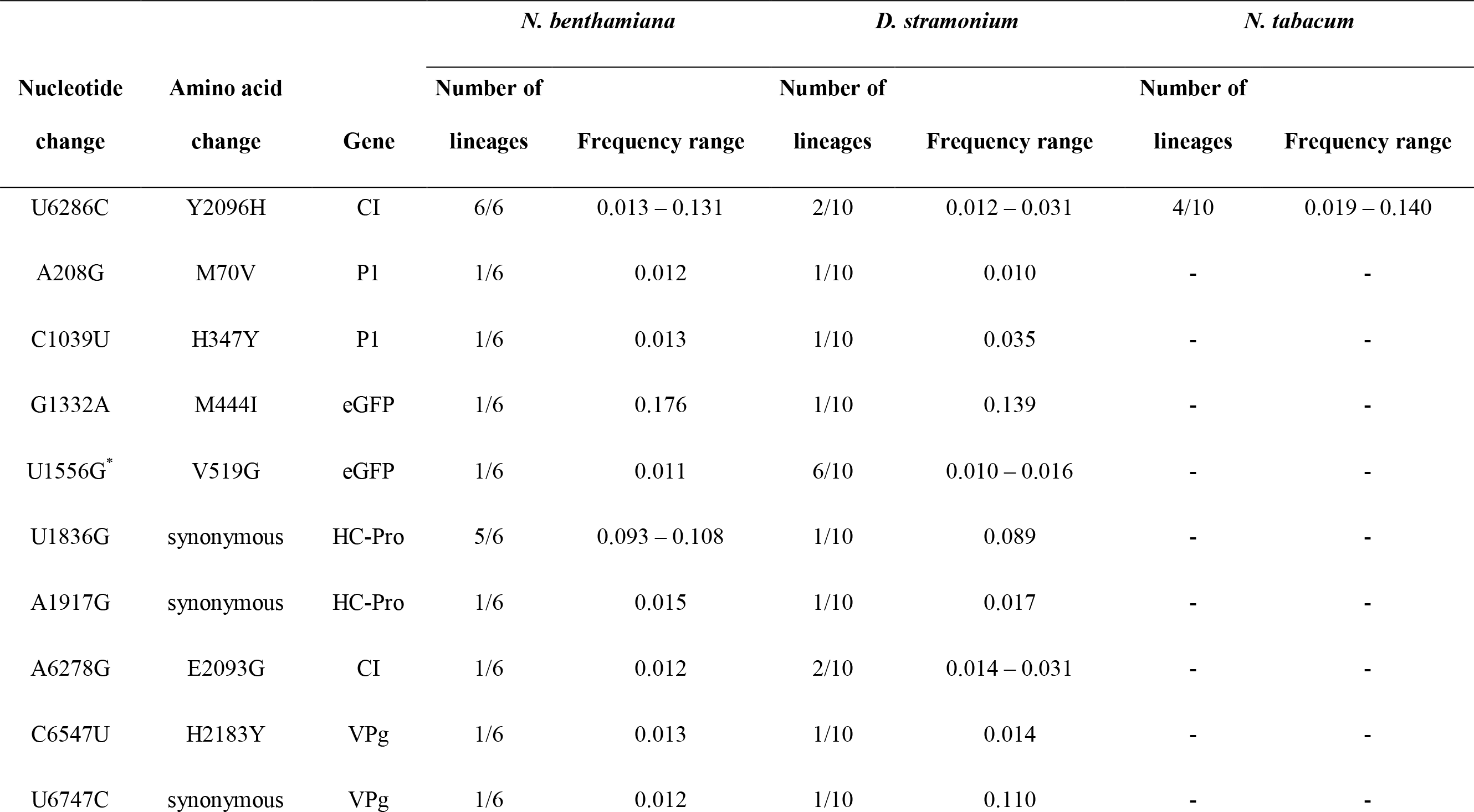
TEV-eGFP mutations shared in the different hosts

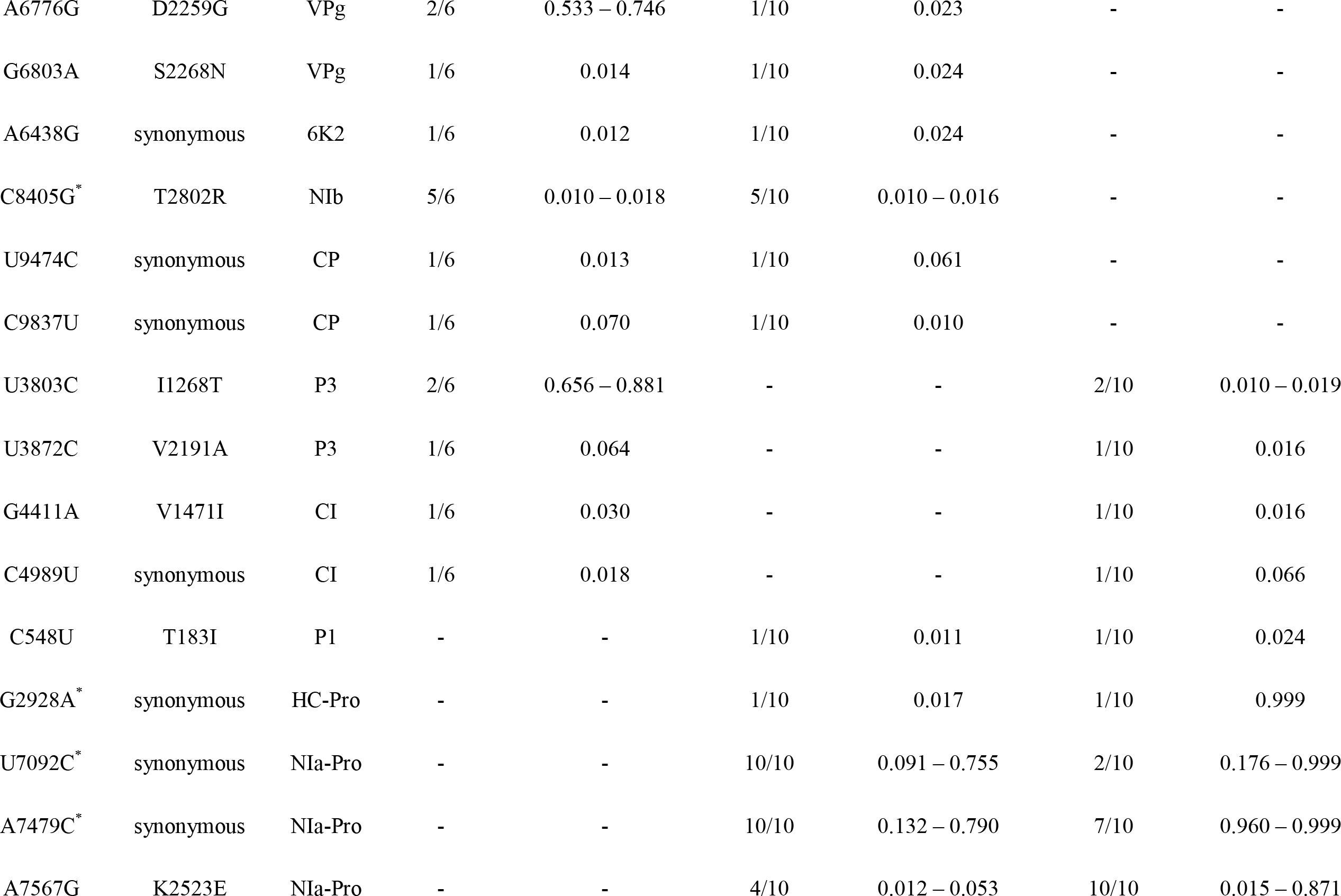

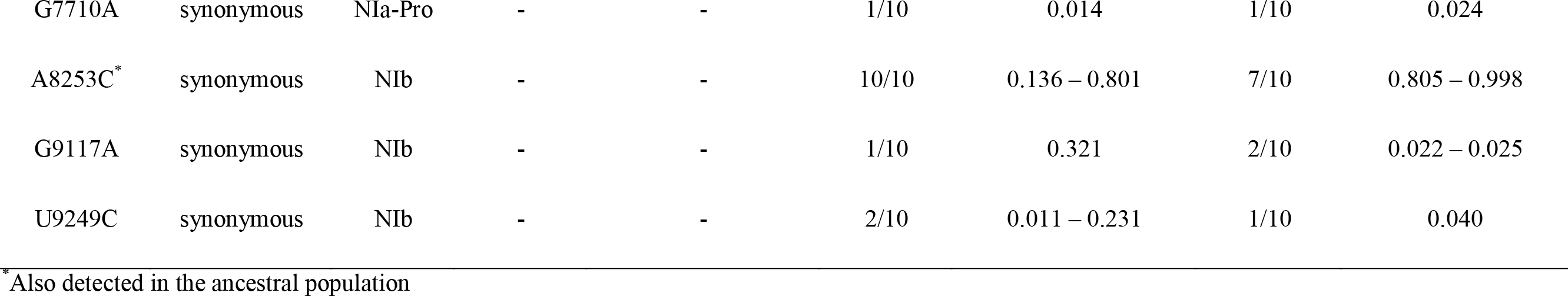

The synonymous mutations U7092C, A7479C and A8253C, that are shared between *D. stramonium* and *N. tabacum*, are present in the highest number of lineages and reach higher frequencies comparing all shared mutations detected in the three hosts. Despite of these mutations already being present in the ancestral population, the frequencies at which these mutations are present display interesting patterns. In both *D. stramonium* and *N. tabacum* the mutations A7479C and A8253C are always detected at the same frequency within each lineage, suggesting a strong linkage between these mutations (Additional file 1: Fig. S1). Furthermore, the U7092C mutation never appears together with the former two mutations (Additional file 1: Fig. S1), suggesting that this mutation occurs in another haplotype and that there may be sign epistasis between these two combinations of synonymous mutations. Interestingly, the ancestral U7092C, A7479C and A8253C mutations were not detected in the *N. benthamiana* lineages, demonstrating the differences in host-pathogen interactions.

Host-specific mutations were mostly found in the evolved TEV-eGFP lineages of *N. benthamiana* (Fig. 3 and Table 2). In this host, a total number of 7 specific mutations were detected, all of them being nonsynonymous. In *D. stramonium* no host-specific mutations were detected. And in *N. tabacum* only one host-specific mutation was detected in the 3’UTR (Table 2). Note that host specific mutations were defined as mutations detected in at least half of the evolved lineages. For more information on the mutations found in the three hosts please see Additional file 2: Tables S1-S3.

**Table 2.** Host specific mutations in the evolved TEV-eGFP lineages.

|  | Nucleotide<br>change | Amino acid<br>change | Gene | Number of<br>lineages | Frequency<br>range |
| --- | --- | --- | --- | --- | --- |
| *N. benthamiana* | G3797A | G1266E | P3 | 3/6 | 0.291 – 0.664 |
|  | G4380U | E1460D | 6K1 | 3/6 | 0.012 – 0.093 |
|  | U4387C | Y1463H | 6K1 | 4/6 | 0.011 – 0.016 |
|  | C4391U | T1464M | 6K1 | 6/6 | 0.041 – 0.138 |
|  | G4397A | S1466N | CI | 6/6 | 0.012 – 0.019 |
|  | A6771U | L2257F | VPg | 4/6 | 0.027 – 0.201 |
|  | G8909U* | W2970L | NIb | 5/6 | 0.026 – 0.042 |
| *D. stramonium* | - | - | - | - | - |
| *N. tabacum* | G10253A |  | 3'UTR | 10/10 | 0.025 – 0.040 |

\*Also detected in the ancestral population.

### Viral accumulation and within-host competitive fitness

We measured virus accumulation 10 dpi, by RT-qPCR for a region within the coat protein gene (*CP*). In both host species, we found no statistically significant differences (*t*-test with Holm-Bonferroni correction) between TEV, TEV-eGFP and the lineages of TEV-eGFP evolved in that host (Fig. 4).

**Figure 4.**
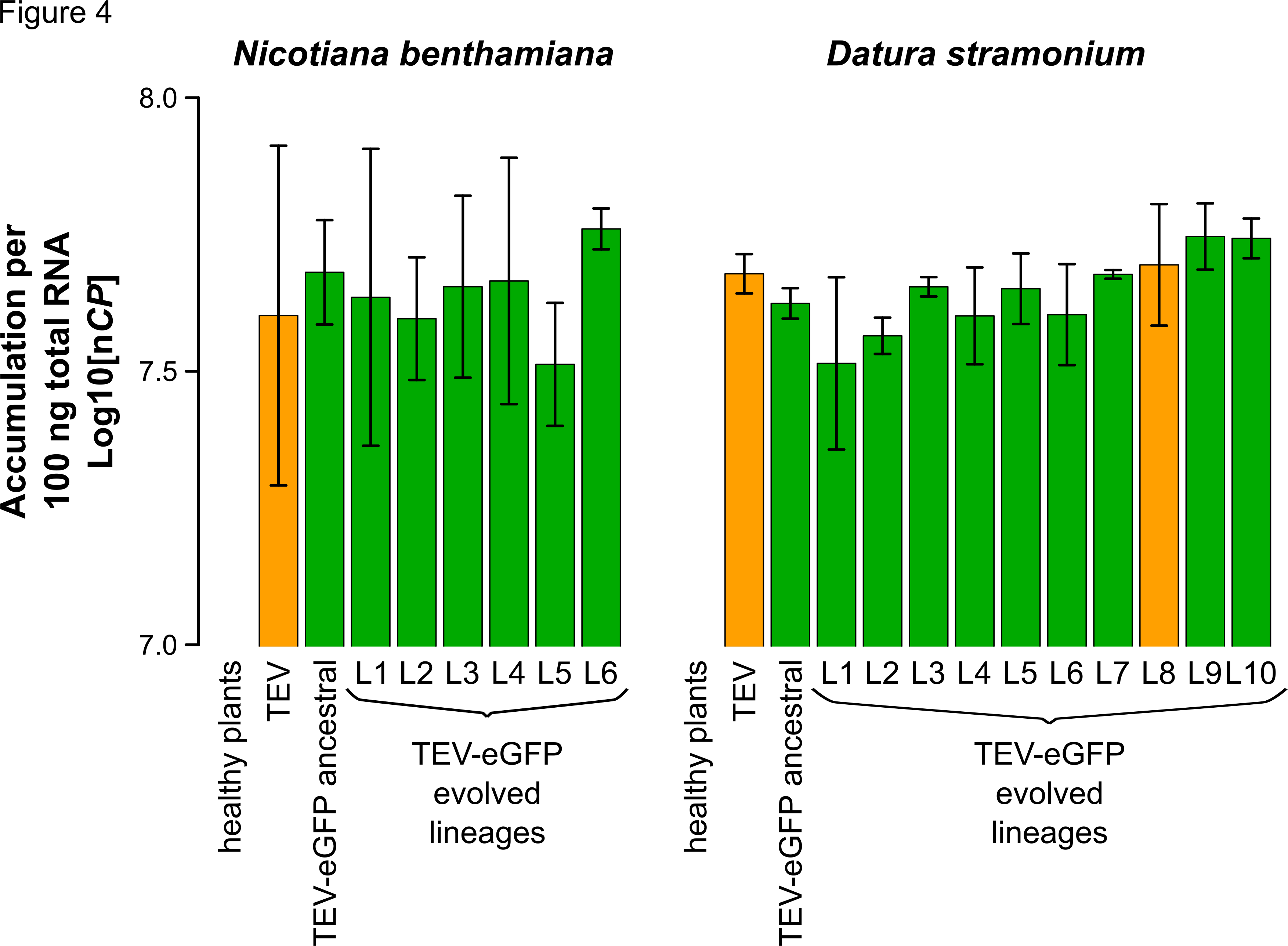
Virus accumulation of the evolved and ancestral lineages. Virus accumulation, as determined by accumulation experiments and RT-qPCR at 10 dpi, of TEV, the ancestral TEV-eGFP, and the evolved TEV-eGFP lineages in the corresponding hosts. TEV and the evolved lineage with a deletion in the *eGFP* gene are indicated with the orange bars. The ancestral TEV-eGFP and the evolved lineages with an intact *eGFP* gene are indicated with the green bars. Error bars represent SD of the plant replicates.

We then measured within-host competitive fitness by means of head-to-head competition experiments with TEV-mCherry, a virus with a different marker but similar fitness to TEV-eGFP [21]. Here we observed interesting differences between TEV and TEV-eGFP in the two different hosts. Whereas the TEV-eGFP had lower fitness than the wild-type virus in *D. stramonium* (Fig. 5, compare TEV and ancestral TEV-eGFP; *t*-test: *t*_4_ = 13.438, *P* < 0.001), there was no difference in *N. benthamiana* (Fig. 5, compare TEV and ancestral TEV-eGFP; *t*-test: *t*_4_ = −1.389, *P* = 0.237). Our results therefore suggest that although there is a fitness cost associated with the *eGFP* cistron in *N. tabacum* [19] and *D. stramonium,* there is none in *N. benthamiana.* Interestingly, *N. tabacum* and *N. benthamiana* are more closely related to each other than either species is to *D. stramonium*, and yet the host species has a strong effect on the costs of a heterologous gene.

**Figure 5.**
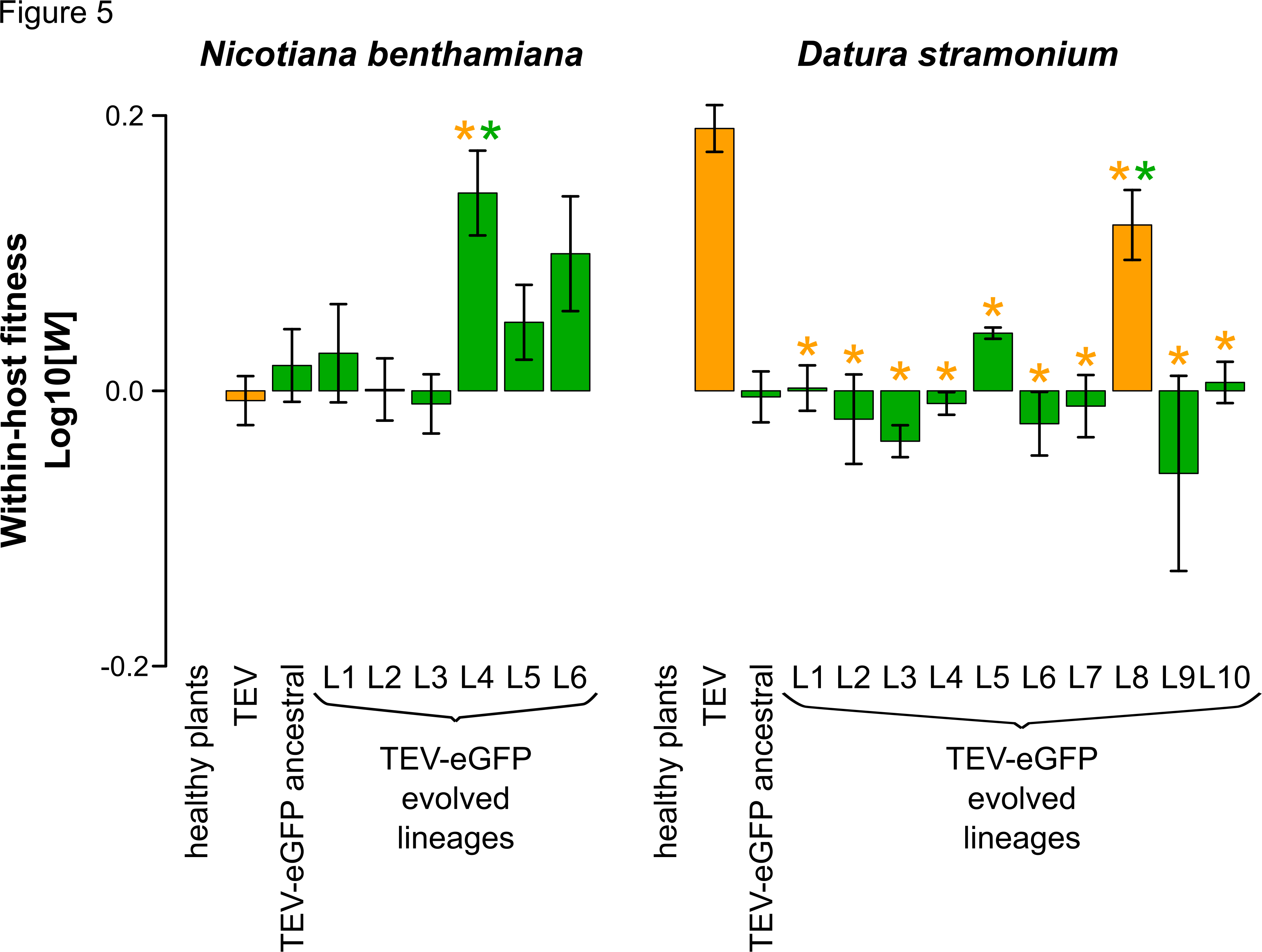
Within-host competitive fitness of the evolved and ancestral lineages. Fitness (*W*), as determined by competition experiments and RT-qPCR of the different viral genotypes with respect to a common competitor; TEV-mCherry. *W* was determined at 10 dpi, of TEV, the ancestral TEV-eGFP, and the evolved TEV-eGFP lineages in the corresponding hosts. TEV and the evolved lineage with a deletion in the *eGFP* gene are indicated with the orange bars. The ancestral TEV-eGFP and the evolved lineages with an intact *eGFP* gene are indicated with the green bars. The orange asterisks indicate statistical significant differences of the evolved lineages as compared to TEV (Mest with Holm-Bonferroni correction). The green asterisks indicate statistical significant differences of the evolved lineages as compared to the ancestral TEV-eGFP (*t*-test with Holm-Bonferroni correction). Error bars represent SD of the plant replicates.Figure 1 TEV-eGFP

For the lineages evolved in *D. stramonium,* only for 1/10 lineages there was a significant increase in competitive fitness compared to the ancestral TEV-eGFP observed (Fig. 5, L8; *t*-test with Holm-Bonferroni correction: *t*_4_ = −6.890, *P* = 0.002). This lineage is the only one to have a deletion in the *eGFP* insert. In *N. benthamiana*, 1/6 lineages had a significant increase in within-host fitness (Fig. 5, L4; *t*-test with Holm-Bonferroni correction: *t*_4_ = −5.349, *P* = 0. 006). However, this increase in fitness probably is not associated with the large genomic deletion for three reasons: (*i*) the wild-type TEV without the *eGFP* cistron has a similar fitness compared to the ancestral TEV-eGFP, suggesting no deletions in *eGFP* would be beneficial, (*ii*) the RT-PCR results show that this variant occurs at a low frequency in the population, and therefore is unlikely to effect strongly the results of the competition assay, and (*iii*) this deletion variant remains at low frequency during the next round of passaging (Fig. 2B), suggesting that while frequency-dependent selection might occur, its fitness is not higher than the coevolving full-length TEV-eGFP. Moreover, another lineage of *N. benthamiana* where we did not detect any deletions, also appeared to have increased in fitness (Fig. 5, L6; *t*-test: *t*_4_ = −4.0792, *P* = 0.015), however, after the Holm-Bonferroni correction not significantly. Interestingly, the lineage that did increase its fitness significantly (L4) is the only lineage that contains mutations in the 6K2 protein in this host (Additional file 2: Table S1). Therefore we speculate that single-nucleotide variation is one of the main driving forces for an increase in TEV-eGFP fitness in *N. benthamiana*.

These fitness measurements show that most lineages failed to adapt to the new host species. However, in the two cases that there were significant fitness increases, the underlying genetic changes were consistent with the expected route of adaptation. In *D. stramonium*, where *eGFP* has a high fitness cost, this sequence was deleted. In *N. benthamiana,* where *eGFP* apparently has not fitness cost, host-specific single-nucleotide variation was observed.

## Discussion

We set out to explore the hypothesis that differences in virulence for different hosts could have an effect on the rate of virus adaptation in each host [13]. Although we find this hypothesis simple and provocative, the observed patterns in our experiments suggest that even in a controlled laboratory environment, reality will often be complex and hard to predict. We used a virus expressing an eGFP fluorescent marker in the hope that the loss of this marker could serve as a real-time indicator of adaptation. However, there were complications with this method, and a loss of fluorescence was only observed in a single *D. stramonium* lineage. RT-PCR and Illumina sequencing confirmed the loss of the eGFP marker in this case, and its integrity in all other lineages. The data of our competitive fitness assay demonstrate why the maker sequence was probably rather stable in *N. benthamiana*; *eGFP* does not appear to have a cost in this host species, in sharp contrast to the strong fitness cost observed in *D. stramonium* as well as previously observed in the more closely related host *N. tabacum* [19]. We expect that the marker will eventually be lost, but only due to genetic drift and therefore at a slow rate.

What mechanisms might underlie the difference in the fitness costs of *eGFP* marker in these two host plants? In a previous study, we showed that the loss of the *eGFP* marker occurred more rapidly as the duration of each passage was increased [19]. During long passages transmission bottlenecks are more spaced on time, and much larger census population sizes are reached. However, there is also much greater scope for virus movement into the newly developing host tissues. As for *N. tabacum* [19], here we again observed that the *eGFP* marker does not affect virus accumulation, whereas it does lower competitive fitness in *D. stramonium.* These observations suggest that the effects of the *eGFP* marker on virus movement are the main reason for selection against the marker. However, marker loss in *D. stramonium* appears to occur much slower compared to *N. tabacum* [19], indicating poor virus adaptation to this alternative host. Given the high virulence of TEV for *N. benthamiana,* including strong stunting, there will be limited virus movement during infection and thus significantly lower population census sizes. Hence, we speculate that in *N. benthamiana* the limited scope for virus movement and accumulation-due to the virus’ virulence itself-might mitigate the cost of the *eGFP* marker. Alternatively, cell-to-cell and systemic virus movement in *N. benthamiana* might be so slow that the addition of the *eGFP* marker matters little. A slow systemic virus movement may also explain why in the second round of infection four lineages failed to re-infect *N. benthamiana*, as initial virus accumulation appeared to be very low until possible virus adaptation by means of point mutations occurred. These results are at odds with our expectations, but they nevertheless have some interesting implications. First, host species changes can apparently ameliorate the costs of exogenous genes. Although strong virus genotype-by-host species interactions have been previously shown for TEV [17], we did not anticipate that a such a simple difference (the presence of *eGFP*) could also be subjected to such an interaction. These results suggest that when considering the evolution of genome architecture, host species might play a very important role, by allowing evolutionary intermediates to be competitive. For example, for TEV we have shown that the evolution of an alternative gene order through duplication of the *NIb* replicase gene is highly unlikely, as this intermediate step leads to significant decreases in fitness, making the trajectory to alternative gene orders inaccessible [34]. If *NIb* duplication has a similar interaction with host species as the *eGFP* insert has, then an alternative host species could act as a stepping-stone and hereby increase the accessibility of the evolutionary trajectory to alternative gene orders. Similar effects of environmental change have been noted in other studies [35]. The generality of these results has not been addressed yet using other viruses with altered genome architecture, but the possibilities are tantalizing. Second, our results could also have implications for assessing the biosafety risks of the genetically modified organisms. Our results suggest that extrapolating fitness results from a permissive host to alternative hosts can be problematic, even when the scope for unexpected interactions appears to be limited, as would be the case for the addition of eGFP expression. In other model systems, unexpected interactions between heterologous genes and host species have also been reported [36].

Our results were not consistent with the hypothesis that high virulence could slow down the rate of adaptation, as in each host only a single lineage had evolved higher fitness. The low rate of adaptation observed was consistent with a previous report [18], although we used passages of a longer duration here and had therefore expected more rapid adaptation [19]. Given the low rate at which lineages adapted in this experiment, however, we do not consider that our results provide strong evidence against the hypothesis. Nevertheless, our results do stress that differences in host biology can have a much stronger effect on evolutionary dynamics than differences in virus-induced virulence between host species. An alternative way to tackling the question of the effects of virulence on adaptation might be to use a biotechnological approach; hosts which have different levels of virulence can be engineered, to ensure the main difference between host treatment is microparasite-induced virulence. For example, plant hosts could be engineered to express antiviral siRNAs at low levels. Such an approach would allow for a more controlled test of the hypothesis suggested here, whilst probably not being representative for natural host populations. On the other hand, such experiments could perhaps help shed light on the effects of virulence on adaptation in agroecosystems or vaccinated populations.

## Conclusions

A host species jump can be a game changer for evolutionary dynamics. A non-functional exogenous sequence –*eGFP*– which is unstable in its typical host, has shown to be more stable in two alternative host species for which TEV has both lower and higher virulence than in the typical host. In addition, *eGFP* does not appear to have any fitness effects in the host for which TEV has high virulence. These observations clashed with the hypothesis that high virulence slows down the rate of adaptation. Moreover, when considering the evolution of genome architecture, host species jumps might play a very important role, by allowing evolutionary intermediates to be competitive.

## Declarations

### Ethics approval and consent to participate

Not applicable

### Consent for publication

Not applicable

### Availability of data and material

The raw read data from Illumina sequencing is available at SRA with accession: SRP075180. The fitness data has been deposited on LabArchives with doi: 10.6070/H4N877TD.

## Author’s contributions

AW, MPZ and SFE designed the study. AW and MPZ performed the experiments. AW, MPZ and SFE analyzed the data and wrote the manuscript. All authors read and approved the final manuscript.

## Acknowledgements

We thank Francisca de la Iglesia and Paula Agudo for excellent technical assistance.

## Additional files

### Additional file 1

**Figure S1.** Frequency of mutations found in both *D. stramonium* and *N. tabacum.* Mutations detected in both *D. stramonium* and *N. tabacum* that were present in all the lineages of either one of these hosts. The frequency of these mutations in either the ancestral population (anc) or the different lineages (L1-L10) is given by the color-coded points. The points are connected by the broken lines to emphasize the trend in the data.

### Additional file 2

**Table S1.** Mutations detected in the *Nicotiana bentiamiana* lineages as compared to the ancestral lineage

**Table S2.** Mutations detected in the *Datura stramonium* lineages as compared to the ancestral lineage

**Table S3.** Mutations detected in the *Nicotiana tabacum* lineages as compared to the ancestral lineage

